# PHIST: fast and accurate prediction of prokaryotic hosts from metagenomic viral sequences

**DOI:** 10.1101/2021.09.06.459169

**Authors:** Andrzej Zielezinski, Sebastian Deorowicz, Adam Gudyś

## Abstract

**Summary:** PHIST (Phage-Host Interaction Search Tool) predicts prokaryotic hosts of viruses from their genomic sequences. It improves host prediction accuracy at species level over current alignment-based tools (on average by 3 percentage points) as well as alignment-free and CRISPR-based tools (by 14–20 percentage points). PHIST is also two orders of magnitude faster than alignment-based tools making it suitable for metagenomics studies.

**Availability and implementation:** GNU-licensed C++ code wrapped in Python API available at: https://github.com/refresh-bio/phist

**Supplementary information:** Supplementary data are available at publisher Web site.

## 1 Introduction

Viruses of prokaryotes constitute the vast majority of the global virosphere and play an important role in balancing ecosystems by regulating the composition of bacteria and archaea worldwide. Novel viruses are routinely discovered by metagenomic studies in a wide range of environments (Paez-Espino *et al*., 2016, Nayfach *et al*., 2021). However, little is known about their biology and the hosts with which they interact since isolation and characterization of novel viruses is remarkably laborious and time-consuming (Edwards *et al*., 2016). Computational tools have been developed to predict prokaryotic hosts from metagenome-derived virus sequences based on the molecular signals of virus-host coevolution, including sequence homology (alignment-based tools), sequence composition similarity between viruses and their hosts (alignment-free tools), and matches to host-encoded CRISPR spacers (Coclet *et al*., 2021).

Alignment-based tools [e.g., BLASTN (Altschul *et al*., 1997), Phirbo (Zielezinski *et al*., 2021)] produce the highest host prediction accuracy of 25–44% at the species level (i.e., percentage of viruses with a correctly predicted host species; for details see Supplementary Material). Nevertheless, these tools have considerable time and memory requirements for large datasets, which hinders their use in metagenomic studies. Alignment-free tools [e.g., WIsH (Galiez *et al*., 2017)] compensate for low speed at the cost of reduced prediction accuracy (18–28% at the species level). Finally, CRISPR-based tools [e.g., SpacePHARER (Zhang *et al*., 2021)] can provide direct evidence supporting virus–host interactions, but these methods have low prediction accuracy because the majority of bacteria lack a CRISPR system. Here, we introduce PHIST, a simple tool that combines benefits of alignment-based, alignment-free, and CRISPR-based approaches to allow a fast and accurate host prediction that can be performed on a standard workstation or even a personal laptop computer.

## 2 Methods

PHIST takes as input two directories containing compressed or uncompressed FASTA files of genomic sequences of viruses and candidate hosts, respectively. The tool links viruses to hosts based on the number of *k*-mers shared between their sequences. PHIST is built upon the Kmer-db tool (Deorowicz *et al*., 2019) using its recently-developed mode of determining numbers of shared *k*-mers (Supplementary Material). PHIST outputs a CSV file with a matrix of common *k*-mer counts between every virus and prokaryote, and a summary CSV file reporting top scoring hosts for each virus. Additionally, PHIST provides *P*-values for the predicted virus-host pairs (Supplementary Material). Information on reference datasets and benchmarking host prediction tools is provided in Supplementary Material.

## 3 Results

We evaluated the performance of PHIST on a reference dataset of 2,288 viral genomes and 62,493 complete and draft prokaryotic genomes from (Wang *et al*., 2020). PHIST correctly identified the hosts species for 29.5% of viruses representing an improvement by 4 percentage points over Phirbo and BLASTN (Fig. 1A). A similar, minor enhancement at the species level (1–2 percentage points) was obtained on two previously published benchmark sets (Edwards *et al*., 2016, Galiez *et al*., 2017) (Supplementary Material). The tool also had the highest host prediction accuracy at the species level when considering the top five predicted host species (i.e., host species with highest similarity to the query virus) (Fig. 1B). PHIST recovered most of the virus-host interaction pairs that were correctly predicted by other approaches. Specifically, our tool recalled 86% (486 out of 561), 85%, 93%, 85%, and 85% virus-host pairs that were correctly predicted by BLASTN, Phirbo, WIsH, SpacePHARER, and PILER-CR, respectively (Fig. 1C).

**Fig. 1.**
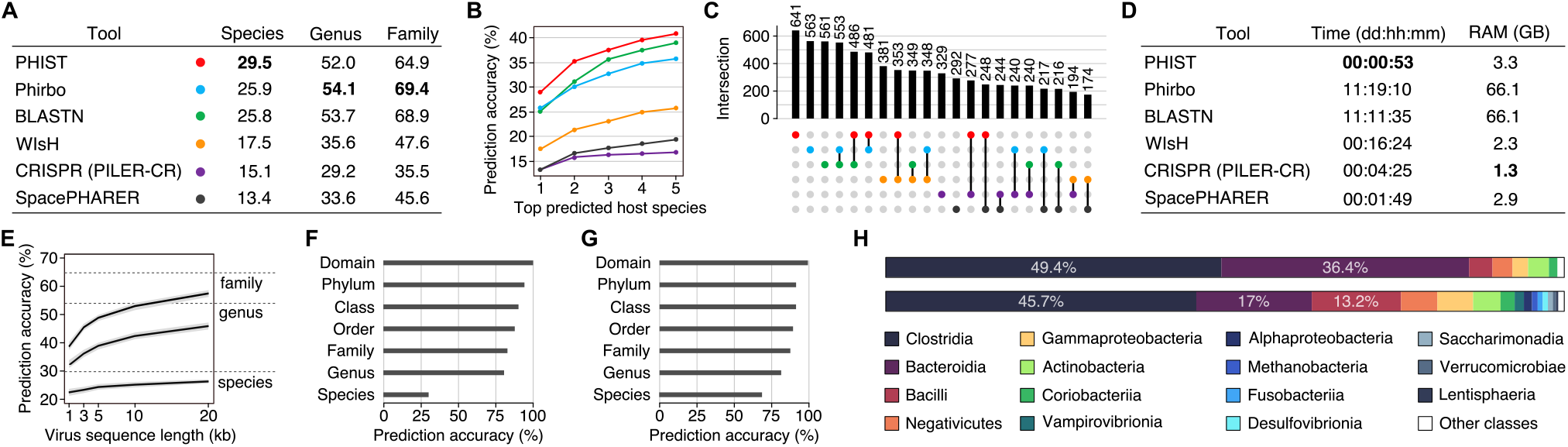
(**A**) Host prediction accuracy for 2,288 viral genomes against 62,493 candidate hosts, binned by host taxonomic level. (**B**) Host prediction accuracy at the species level across five highest scoring prokaryotic species for each virus sequence (the prediction was scored as correct if the correct host species was among the first five predicted hosts). (**C**) Unique and shared correct virus-host predictions among the tools. The bar chart indicates the intersection size of viruses with correct host prediction at the species level. Connected dots on the bottom panel indicate which pair of tools is considered for each intersection. (**D**) Runtime and memory usage for 2,288 viruses and 62,493 prokaryotes. Time and memory measurements of Phirbo also include BLASTN, and SpacePHARER includes PILER-CR analysis. (**E**) Prediction accuracies for contigs subsampled at various lengths from the 2,288 virus genomes. The solid line shows the mean prediction accuracy at host taxonomic level and the grey shade indicates the 95% confidence interval. The horizontal dashed line marks the prediction accuracy using the full-length viral genomes. (**F, G**) Proportions of congruent predictions for viral contigs between PHIST and those in Paez-Espino *et al*. and Nayfach *et al*., respectively. (**H**) Viral assignments to human gut bacteria and archaea. Distribution of host classes reported by Nayfach *et al*. (upper bar chart) and a set of PHIST predictions on viruses that were not assigned to hosts by Nayfach *et al* (lower bar chart).

PHIST took an hour to process the dataset (approximately 143 million pairwise comparisons between 2,288 viruses and 62,492 candidate hosts) on a 16-core 2.1GHz Intel Xeon, almost 300 times faster than BLASTN and Phirbo, 16 times faster than WIsH, and 2-4 times faster than CRISPR-based tools (Fig. 1D).

To evaluate the performance of host prediction for short viral contigs, we randomly subsampled fragments of different lengths (1, 2, 5, 10, and 20 kb) from each of the 2,288 viral genomes. The mean host prediction accuracy at the genus and family levels dropped by 9 and 11 percentage points, respectively, for 20 kb to 3 kb long contigs (Fig. 1E). At the species level, however, the prediction accuracy fell by only 3 percentage points (from 26% to 23%) for 20 kb to 3 kb long contigs. We also tested PHIST on a set of 125,842 metagenomic viral contigs (MVCs) of 11 kb median length obtained from various environments (Paez-Espino *et al*., 2016). The original host prediction used CRISPR-spacer and tRNA sequence matches, and assigned hosts for only 7.9% of the MVCs. PHIST annotated 99% of the MVCs; moreover, the predictions matched the original predictions in 30% at the species level and achieved 80% consistency at the genus level (Fig. 1F).

The resulting accuracies can be considered as lower bounds since metagenomic studies restrict the set of candidate host genomes to those present in the sample (Galiez *et al*., 2017). Thus, we used a set of 189,680 metagenomic viral sequences from human gut provided by Nayfach *et al*. (2021), who predicted the hosts among 286,997 genomes of bacteria and archaea from the Unified Human Gastrointestinal Genome (UHGG) collection (Almeida *et al*., 2020). The original host prediction used a combination of CRISPR-spacer matches and whole-genome alignments between viruses and candidate hosts, (Nayfach *et al*., 2021) and assigned hosts to 90% of the viral contigs (*n* = 170,072). PHIST annotated 99.95% of the viral contigs (*n* = 189,586) and provided an overall 81% agreement on host taxonomy with the original predictions (Supplementary Material). Using the Nayfach *et al*. (2021) annotation as reference, PHIST obtained 68% and 81% host prediction accuracy at the species and genus levels, respectively (Fig. 1G). The remaining viral genomes that were not assigned to hosts in Nayfach *et al*. (*n* = 19,514) were connected by PHIST mostly to Clostridia and Bacteroidia (Fig. 1H, lower bar chart), which were also two dominant classes of bacteria in Nayfach *et al*. predictions (Fig. 1H, upper bar chart). Computing all ∼54 billion pairwise scores between 189,680 viruses and 286,997 prokaryotes required just 3.5 hours and 25 GB of RAM (Supplementary Material).

## 4 Conclusion

PHIST predicts hosts species with higher accuracy than alignment-based tools, and allows for rapid analysis of large-scale genomic and metagenomic datasets on non-specialized hardware.

## Data availability

All data analyzed in this study are available from previously published studies (Supplementary Material).

## References

Almeida, A. et al. (2020) A unified catalog of 204,938 reference genomes from the human gut microbiome, Nat. Biotechnol., 39, 105–114.

Altschul, A.F. et al. (1997) Gapped BLAST and PSI-BLAST: a new generation of protein database search programs, Nucleic Acids Res., 25, 3389–3402.

Coclet, C. and Roux, S. (2021) Global overview and major challenges of host prediction methods for uncultivated phages, Curr. Opin. Virol., 49, 117–126.

Deorowicz, S. et al. (2019) Kmer-db: instant evolutionary distance estimation, Bioinformatics, 35, 133–136.

Edwards, R. et al. (2016) Computational approaches to predict bacteriophage–host relationships, FEMS Microbiol. Rev., 40, 258–272.

Galiez, C. et al. (2017) WIsH: who is the host? Predicting prokaryotic hosts from metagenomic phage contigs, Bioinformatics, 33, 3113–3114.

Nayfach, S. et al. (2021) Metagenomic compendium of 189,680 DNA viruses from the human gut microbiome. Nat. Microbiol., 6, 960–970.

Paez-Espino, D. et al. (2016) Uncovering Earth’s virome. Nature, 536, 425–430.

Wang, W. et al. (2020) A network-based integrated framework for predicting virus–prokaryote interactions. NAR Genom. Bioinform., 2, 1–19.

Zhang, R. et al. (2021) SpacePHARER: Sensitive identification of phages from CRI-SPR spacers in prokaryotic hosts, Bioinformatics, Published on-line: 01 April 2021.

Zielezinski, A., Barylski, J., Karlowski, W.M. (2021) Taxonomy-aware, sequence similarity ranking reliably predicts phage-host relationships. BMC Biol., in press.

