## Supplementary material for "PHIST: fast and accurate prediction of prokaryotic hosts from metagenomic viral sequences"

Supplementary Information

Andrzej Zielezinski, Sebastian Deorowicz, Adam Gudys

September 6, 2021

### 1 PHIST description

#### 1.1 The algorithm

PHIST (Phage-Host Interaction Search Tool) predicts prokaryotic hosts of viruses from their genomic sequences. The tool calculates the number of shared $k$-mers between all-*versus*-all viral and candidate host sequences. By default, the tool uses $k = 25$ as this length is considered the lower boundary of the size range of mechanisms leading to the retention of identical sequences in the genomes of a virus and its host (i.e., CRISPR spacers, recombination sites) (?). Similar to other $k$-mer-based tools, PHIST operates on *canonical* $k$-mers (i.e., without differentiation between $k$-mers and their reverse complement).

PHIST takes as input two directories containing FASTA files (gzipped or not) with genomic sequences of viruses and candidate hosts, respectively. First, PHIST runs Kmer-db (?) to build a database of all $k$-mers present in a query set of viral genomic sequences. Then, PHIST runs Kmer-db with the *new2all* mode to count common $k$-mers between the query prokaryotic genomes and all the viruses in the database. Finally, for each virus PHIST returns prokaryote(s) with the highest number of $k$-mers shared with the query viral sequence.

#### 1.2 *P*-values

For a predicted pair of virus $V$ and host sequences $H$ of lengths $m$ and $n$ respectively, PHIST provides probability $P$ of finding at least $\ell$ common $k$-mers of length $k$ between $V$ and $H$ under the null hypothesis that the nucleotide composition of both genomes is independent. Notably, PHIST provides the upper bound on $P$-value assuming the common $k$-mers overlap in the virus and host genome by $k-1$ nucleotides, thus encompassing a single sequence of length $s = k+\ell-1$. The number of all possible canonical sequences of length $s$ is given by $N = 4^s/2$ if $s$ is odd or $(4^s + 4^{s/2})/2$ if $s$ is even. Then, the expected number of common sequences of length $s$ between two genomes of lengths $m$ and $n$ is given by:

$$\lambda^s_{m,n} = \frac{(m - s + 1) \times (n - s + 1)}{N}. \tag{1}$$

The probability of finding at least one common sequence of length $s$ in viral and host genomes can be approximated using the Poisson cumulative distribution function:

$$P(i \geq 1) = 1 - P(i = 0) = 1 - exp(-\lambda^s_{m,n}). \tag{2}$$

Since $P$-value calculated by Eq. 2 only describes the significance of a single virus-host comparison, PHIST also provides $P$-value adjusted by the multi-testing Bonferroni correction (i.e., $P$-value is multiplied by the number of prokaryotes being input). If the adjusted $P$-value ended up $> 1.0$, it would be rounded down to 1.0.

### 2 Datasets

The sequence data of viruses and prokaryotes as well as the information on reference virus-host assignments were retrieved from five previously published studies (Table 1).

Table 1: Reference datasets used in this study.

| Dataset | #Viruses | #Prokaryotes | Size [GB] | Reference |
|---|---|---|---|---|
| Wang *et al.* (2021) | 2,288 | 62,492 | 250 | (?) |
| Edwards *et al.* (2016) | 820 | 2,699 | 8 | (?) |
| Galiez *et al.* (2017) | 1,420 | 3,780 | 13 | (?) |
| Paez-Espino *et al.* (2016) | 125,842 | 62,492 | 252 | (?) |
| Nayfach *et al.* (2021) | 189,680 | 286,997 | 667 | (??) |

### 3 Host prediction tools

Host prediction tools used in this study were run separately on the **?**, **?**, and **?** datasets. All the tools were run with their default parameters.

#### 3.1 BLASTN

blastn from the BLAST+ package (v.2.7.1) (**?**) was run to query each viral sequence against a database of candidate host genomes.

```
$ cat host/fasta/*.fasta > hosts.fasta |
makeblastdb -in hosts.fasta -dbtype nucl

$ ls virus/fasta/*.fasta | xargs -I {}
blastn -task blastn -query {} -db hosts.fasta -outfmt 6
-num_threads 16 -out blastn/{}.txt
```

#### 3.2 Phirbo

Phirbo (v.1.0) was run based on the ranked lists for virus and prokaryote genomes. Ranked lists for viruses were created directly from the BLASTN output, and the ranked lists for prokaryotes were created based on the Mash (v.2.1) (**?**) distances between all-versus-all prokaryote genomes. Mash was run with default parameters ($k$ = 21, sketch size = 1,000).

```
$ ./phirbo.py --t 16 virus/ host/ phirbo/predictions.csv
```

#### 3.3 WIsH

WIsH (v.1.0) was run with default parameters.

```
$ mkdir modelDir
$ WIsH -c build -g host/fasta/ -m modelDir -t 16
$ mkdir wish
$ WIsH -c predict -g virus/fasta/ -m modelDir -r wish -b -t 16
```

#### 3.4 PILER-CR

PILER-CR (v.1.06) (**?**) was run against all candidate host genomes to extract CRISPR spacer sequences, which were aligned to the viral genomes using BLASTN. BLASTN search was adapted for CRISPR spacer sequences as suggested by **?** (i.e., using the blastn-short task, e-value threshold of 1, a gap open penalty 10, a gap extension penalty 2, a mismatch penalty 1, and a word size 7). Exact or near-exact CRISPR spacer alignments to viral genomes (up to 2 mismatches) were selected. The prokaryotic genomes sharing the highest number of the common hits with viral genome were reported for each virus.

#### 3.5 SpacePHARER

SpacePHARER (Release 5-c2e680a) (**?**) was run based on the CRISPR spacer sequences returned by PILER-CR. For each virus the predicted host with the highest combined score ($S_{comb}$) was reported.

```
$ spacepharer createsetdb virus/fasta/*.fasta targetSetDB
temp/ --threads 16

$ spacepharer createsetdb virus/fasta/*.fasta targetSetDB_rev
temp/ --reverse-fragments 1 --threads 16
```

```
$ spacepharer parsespacer pilercr/*.txtqueryDB --threads 16 -v 0

$ spacepharer createsetdb queryDB querySetDB temp/
--extractorf-spacer 1 --threads 16

$ spacepharer predictmatch querySetDB targetSetDB
targetSetDB_rev spacepharer/predictions.tsv temp/ --threads 16
```

##### 3.6 PHIST

PHIST (v.1.0) was run with default parameters (i.e., $k = 25$).

```
$ ./phist.py --t 16 virus/fasta/ host/fasta
```

#### 4 Benchmark details

Following previous studies (**????**), host prediction tools were evaluated by selecting a top-scored prokaryotic sequence for each virus. Host prediction accuracy is defined as the percentage of viruses whose predicted hosts have the same taxonomic affiliation as their respective known hosts. In case multiple top-scoring hosts are present for a given virus we used two approaches to calculate host prediction accuracy. First approach was used by the previous studies and score a prediction as correct if the true host is among the predicted hosts. In second approach we randomly selected a single host from multiple top-scoring hosts. Tables 2-4 show host prediction accuracies calculated by the two approaches on the **?**, **?**, and **?** datasets, respectively.

Althoug PHIST was outperformed by alignment-based methods (i.e., Phirbo and BLAST) at the genus and family levels, our tool obtained the highest host prediction accuracy (29.5–44.6%) at the species level across all the three datasets.

Table 2: Host prediction accuracies (%) for phage and bacteria genomes from the datasets by Wang *et al.* (2020). Host prediction accuracy calculated by randomly selecting a single host from multiple top-scoring prokaryotes is shown in parentheses. The highest accuracies among the tools for each taxonomic level are in bold.

| **Tool** | **Species (%)** | **Genus (%)** | **Family (%)** |
|---|---|---|---|
| PHIST | **29.5 (29.0)** | 52.0 (51.2) | 64.9 (63.6) |
| Phirbo | 25.9 (25.8) | **54.1 (54.0)** | **69.4 (69.4)** |
| BLASTN | 25.8 (24.9) | 53.7 (53.5) | 68.9 (68.7) |
| WIsH | 17.5 (17.5) | 35.6 (35.6) | 47.6 (47.6) |
| CRISPR (PILER-CR) | 15.1 (13.4) | 29.2 (25.8) | 35.5 (31.4) |
| SpacePHARER | 13.4 (13.4) | 33.6 (33.6) | 45.6 (45.6) |

##### 4.1 Viral contigs subsampled from full-length genomes

Analogously to **?** we evaluated the performance of host prediction for short viral contigs. Specifically, we randomly subsampled fragments of fixed lengths (i.e., 1, 2, 5, 10, and 20 kb) from each of the 2,288 viral genomes. For a given viral genome and a fixed contig length, we randomly chose a sequence segment of that length. If the fixed length was longer than the size of the complete viral genome, we took the entire genome. This procedure was repeated 30 times for each contig length. Hosts were predicted on each subsampling replicate from among 62,492 prokaryotic genomes. Host prediction accuracy was averaged across prediction accuracies of repliactes within each contig length. A 95% confidence interval for the mean host prediction accuracy was estimated using normal distribution.

Table 3: Host prediction accuracies (%) for phage and bacteria genomes from the datasets by ?. Host prediction accuracy calculated by randomly selecting a single host from multiple top-scoring prokaryotes is shown in parentheses. The highest accuracies among the tools for each taxonomic level are in bold.

| **Tool** | **Species (%)** | **Genus (%)** | **Family (%)** |
|---|---|---|---|
| PHIST | **44.6 (43.9)** | 57.6 (56.2) | 66.2 (64.6) |
| Phirbo | 43.2 (43.2) | 58.8 (**58.8**) | 70.5 (**70.5**) |
| BLASTN | 43.2 (42.4) | **58.9** (57.9) | **70.7** (70.2) |
| WIsH | 28.3 (28.3) | 44.3 (44.3) | 49.9 (49.9) |
| CRISPR (PILER-CR) | 15.9 (15.4) | 17.8 (17.4) | 21.6 (21.5) |
| SpacePHARER | 22.3 (21.5) | 25.6 (24.5) | 31.6 (30.9) |

Table 4: Host prediction accuracies (%) for phage and host genomes from the datasets by ?. Host prediction accuracy calculated by randomly selecting a single host from multiple top-scoring prokaryotes is shown in parentheses. The highest accuracies among the tools for each taxonomic level are in bold.

| **Tool** | **Species (%)** | **Genus (%)** | **Family (%)** |
|---|---|---|---|
| PHIST | **33.2 (32.3)** | 48.6 (47.0) | 54.9 (53.2) |
| Phirbo | 31.2 (31.2) | 52.9 (**52.9**) | 61.5 (**61.5**) |
| BLASTN | 31.1 (31.1) | **53.1** (52.7) | **61.7** (61.4) |
| WIsH | 20.6 (20.6) | 43.7 (43.7) | 48.0 (48.0) |
| CRISPR (PILER-CR) | 8.4 ( 8.0) | 12.1 (11.5) | 14.3 (13.6) |
| SpacePHARER | 9.6 ( 9.5) | 18.9 (18.9) | 23.5 (23.5) |

#### 4.2 Metagenomic viral contigs

? described over 3,000 geographically diverse metagenomic samples and identified 125,842 metagenomic viral contigs (MVCc) of median length 11 kb. The original host prediction (?) involved two approaches: matching MVCs to a database of 3.5 million CRISPR spacers found in prokaryotic genomes, and annotating tRNA sequences in corresponding hosts. Peaz-Espino *et al.* assigned 7.9% of MVCs ($n$ = 9,992) to hosts at various taxonomic levels (Supplementary Table S19 in ?). PHIST was run for all 125,842 MVCs using 62,492 prokaryotic genomes from ?. Of note, only the host annotations from Peaz-Espino *et al.* whose taxonomic levels were present in our prokaryotic genomes were kept as the reference host assignments, resulting in 7.4% of MVCs ($n$ = 9,255).

#### 4.3 Human gut microbiome

? provided a set of 189,680 metagenomic viral genomes (MVGs) from 11,810 human gut metagenomes (?). The original host prediction (?) used a combination of CRISPR-spacer matches and $\geq$ 1 kb genome sequence matches to associate MVGs to bacterial and archaeal genomes from the the Unified Human Gastrointestinal Genome (UHGG) collection (?). UHGG contains 286,977 genomes, representing 4,644 bacterial and archaeal species from the human gut that are taxonomically annotated using GTDB-tk v.0.3.1 (GTDB release 89) (?). Following the original host prediction procedure (?), we removed 2,043,531 contigs from UHGG genomes (out of 59,281,966 contigs) that were most likely erroneously assembled (i.e., host region comprised $<$ 50% of the contig length). Nayfach *et al.* assigned 90% of MVGs ($n$ = 170,072) to hosts at different taxonomic levels (Table 4). For example, for 73,141 MVGs host was annotated at the species level while for 35,967 MVGs the lowest host taxonomic level was genus.

PHIST predictions were generally consistent with the host annotations from ?. Specifically, for 81% of MVGs ($n$ = 137,104) the taxonomic affiliation of the predicted hosts matched the taxonomic affiliation provided by Nayfach *et al.* at the lowest host taxonomic rank. Host pre-

Table 5: Host annotation of 170,072 mVGs obtained from **?**

| **Host taxonomic level** | **#MVGs** |
|---|---|
| Species | 73,141 |
| Genus | 35,967 |
| Family | 35,268 |
| Order | 17,716 |
| Class | 5,548 |
| Phylum | 37 |
| Domain | 2,395 |

diction accuracies obtained by PHIST across different taxonomic levels are provided in Table 6.

Table 6: Host prediction accuracies (%) of PHIST on the **?** dataset

| **Host taxonomic level** | **Host prediction accuracy** (%) |
|---|---|
| Species | 68.4 |
| Genus | 80.9 |
| Family | 87.2 |
| Order | 88.8 |
| Class | 90.8 |
| Phylum | 90.8 |
| Domain | 99.9 |

#### 4.4 Runtime and memory usage

Running time and peak memory were measured with /usr/bin/time -v on a Linux PC with 16-core 2.1 GHz Intel Xeon Gold 6130 processor.

Table 7: PHIST runtime and memory usage across the reference datasets

| **Dataset** | **#Viruses** | **#Prokaryotes** | **#Comparisons** | **Time (hh:mm)** | **RAM (GB)** |
|---|---|---|---|---|---|
| Edwards *et al.* | 820 | 2,699 | 2,213,180 | 00:02 | 2.1 |
| Galiez *et al.* | 1,420 | 3,780 | 5,367,600 | 00:03 | 2.6 |
| Wang *et al.* | 2,288 | 62,492 | 142,981,696 | 00:53 | 3.3 |
| Paez-Espino *et al.* | 125,842 | 62,492 | 7,864,118,264 | 01:06 | 21.8 |
| Nayfach *et al.* | 189,680 | 286,997 | 54,437,590,960 | 03:18 | 25.1 |
